# AnnoView enables large-scale analysis, comparison, and visualization of microbial gene neighborhoods

**DOI:** 10.1101/2024.01.15.575735

**Authors:** Xin Wei, Huagang Tan, Briallen Lobb, William Zhen, Zijing Wu, Donovan H. Parks, Josh D. Neufeld, Gabriel Moreno-Hagelsieb, Andrew C. Doxey

## Abstract

The analysis and comparison of gene neighborhoods is a powerful approach for exploring microbial genome structure, function, and evolution. Although numerous tools exist for genome visualization and comparison, genome exploration across large genomic databases or user-generated datasets remains a challenge. Here, we introduce AnnoView, a web server designed for interactive exploration of gene neighborhoods across the bacterial and archaeal tree of life. Our server offers users the ability to identify, compare, and visualize gene neighborhoods of interest from 30,238 bacterial genomes and 1,672 archaeal genomes, through integration with the comprehensive GTDB and AnnoTree databases. Identified gene neighborhoods can be visualized using pre-computed functional annotations from different sources such as KEGG, Pfam, and TIGRFAM, or clustered based on similarity. Alternatively, users can upload and explore their own custom genomic datasets in GBK, GFF, or CSV format, or use AnnoView as a genome browser for relatively small genomes (e.g., viruses and plasmids). Ultimately, we anticipate that AnnoView will catalyze biological discovery by enabling user-friendly search, comparison, and visualization of genomic data. AnnoView is available at http://annoview.uwaterloo.ca

## Background

Gene neighborhoods are regions of genomes containing sets of co-localized genes. For bacteria and archaea, the organization of genes into clusters or operons can provide insight into gene function, regulation, and evolutionary relationships [1–3]. The identification of specialized gene clusters within genomes can help identify diverse physiological and functional traits including biosynthetic gene clusters [4, 5], transport and uptake systems [6], and antibiotic resistance and virulence traits [7–10]. Comparison of orthologous gene neighborhoods can reveal functional diversification [7, 11, 12], identify cases of horizontal transfer and genomic rearrangements [13–15], predict functional associations between genes [2, 16, 17], and infer operons and pathways [18, 19].

There are multiple strategies for analyzing sets of neighboring genes and their patterns of co-localization across genomes. Tools such as Artemis / Artemis Comparison Tool (ACT) [20], IGV [21], Mauve [22], or Jbrowse [23] can be used for pairwise or small-scale gene neighborhood comparison involving one or a few species. These tools are valuable for visualizing and comparing genomic regions of interest and their associated functional annotations. Synteny mapping tools can also aid in the comparison of gene neighborhoods [24–26]. Additionally, public databases and comparative genomics platforms can provide gene neighborhood visualization features, which include the NCBI [27], UCSC [28], STRING[29], and Ensembl databases [30]. For more advanced gene neighborhood analysis, including comparative analysis across larger sets of genomes, several tools and portals have been developed [31–42]. These can provide a more detailed and focused view on gene neighborhoods than genome browsers, facilitate comparison across species in pre-computed databases, and exploration of predicted gene functions from different annotation methods.

Despite existing tools for gene neighborhood comparison, there is still a need for a platform that: 1) is intuitive and easy to use; 2) contains a wide diversity of genomes across the full bacterial and archaeal tree of life; 3) is taxonomically and functional annotated in a comprehensive and consistent manner; 4) is not restricted to a single reference database and allows users to upload their own custom datasets; 5) is capable of comparing relatively large numbers of genomic regions simultaneously; 6) is capable of exploring “unannotated” genes and proteins of unknown function [43]. In order to address the need for a platform with such functionality, here we introduce AnnoView, a versatile and easy-to-use web platform for microbial gene neighborhood exploration. AnnoView enables large-scale gene neighborhood search and visualization across 30,238 bacterial genomes and 1,672 archaeal genomes. Through integration with AnnoTree [44] and the Genome Taxonomy Database (GTDB) [45], AnnoView allows users to identify and explore gene neighborhoods of interest across large sets of genomes, and cluster and visualize them based on gene function composition, with a choice of different annotation methods including KEGG [46, 47], PFAM, and TIGRAM [48]. Additionally, AnnoView allows users to upload custom genome datasets, such as sets of gene neighborhoods from different species, metagenomic contigs, small bacterial genomes, viral genomes, and plasmids. Users can easily share their AnnoView sessions using session URL links or download their data for offline editing.

## Methods

### Construction of the AnnoView database

The AnnoView database was constructed using information from the AnnoTree R95 database and Genome Taxonomy Database (R95) [44, 45]. Gene and functional annotations are derived from the gene prediction and annotation pipeline described previously [44]. In brief, Prodigal [49] with default parameters was used for gene calling from whole genomic FASTA files, resulting in GBK/GFF files and predicted protein-coding sequences for all organisms. Both Pfam and TIGRFAM annotation were performed on all protein sequences using HMMER [50] as well as pfam_scan.pl (ftp://ftp.ebi.ac.uk/pub/databases/Pfam/Tools/) for Pfam domains specifically. For KEGG annotation, DIAMOND [51] was used to identify homologous sequence clusters within the UniRef100 database [52] and existing annotations were transferred. DIAMOND was run with the following parameters: *E*-value ≤ 1e-5, identity ≥ 30%, and query-to-subject and subject-to-query alignment coverage ≥ 70%. This pipeline was used to functionally annotate the genes from 30,238 bacterial genomes and 1,672 archaeal genomes.

For the AnnoView database, short names of and descriptions of functional annotations were retrieved from the above KEGG, Pfam, and TIGRFAM annotation data. Gene coordinates and orientations were extracted from the Prodigal-generated GFF files. An AnnoView SQL database was constructed to store predicted protein sequences, gene functional annotation data (i.e., KEGG, Pfam and TIGRFAM), gene coordinates and strand information (i.e., + or −), CDS length, as well as GTDB taxonomy information. All protein sequences in the AnnoView database were then used to build a DIAMOND (2.1.8) database [51] that can be searched to find similar proteins to a query. Once DIAMOND searches are performed by the user, the backend server will run MySQL queries on all sequence hits to identify proteins within a ±10 kb window size. Protein-coding genes that span the edges of this window are excluded. This process is highly efficient due to the presence of pre-computed data and indexes in the MySQL database, allowing for quick execution of MySQL queries.

### Web server implementation

The AnnoView web server adopts a client-server website design (**Supplementary Figure 1**). For the frontend client, we used react.js to build the user interface (https://reactjs.org/) and d3.js (https://d3js.org) for visualizing gene neighborhoods. The Python-based Flask web framework (https://flask.palletsprojects.com) was used for backend server development. Pre-computed gene neighborhood data are stored in a MySQL (https://www.mysql.com/) database. Gene neighborhood data resulting from user queries and user-uploaded data are stored in a MongoDB (https://www.mongodb.com/) database. The pseudocode for the algorithm that clusters gene neighborhoods based on a gene of interest (**Supplementary Figure 2**) is a modified version of the Smith-Waterman algorithm for local sequence alignment [53] applied to vectors of gene lists where each gene is labeled by its annotation.

### Upload workflow for NCBI data retrieval and gene annotations

In addition to retrieval of gene neighborhood information from the AnnoTree [44] and GTDB [45] database, a bioinformatic workflow was outlined for users to retrieve gene neighborhood information directly from the NCBI database, which includes additional genomic information not available in the GTDB. The workflow involves: (1) retrieval of gene neighborhood datasets in GBK format from NCBI; (2) annotation of gene neighborhoods with taxonomic information; (3) KEGG and Pfam functional annotation of gene neighborhoods; (4) sorting of gene neighborhoods by selection of a gene/protein of interest as the “center gene”; and (5) visualization of gene neighborhoods in AnnoView. The detailed NCBI-based workflow and associated code is accessible at the following GitHub repository: https://github.com/satellite830/AnnoView

## Results

### The AnnoView platform

AnnoView is a gene neighborhood browser for microbial genomes available at http://annoview.uwaterloo.ca/. There are two modules in AnnoView: (1) the “upload” module allows users to upload genomic neighborhoods in GBK, GFF, and CSV format; and (2) the “GTDB search” module allows users to query the GTDB/AnnoView genome database for matches to an input protein sequence. After a protein query is searched against the AnnoView database, the server returns genomic neighborhoods surrounding identified matches (genes corresponding to protein homologs). Pre-computed functional annotations in KEGG, Pfam, and TIGRFAM are then available to the user for further exploration and visualization. The protein homology search criteria includes *E*-value, coverage cut-off, and a choice of database to search (bacteria/archaea).

After the homology search, the user can further select a subset of protein hits for genomic neighborhood display by taxonomy or by *E*-value on an “intermediate” results page. This intermediary page solves potential issues of overwhelming number of sequence matches, which need further filtering. The “Select by taxonomy” option is intended for a user interested in exploring genomic neighborhoods in a specific lineage, whereas “Select by *E*-value” is intended for users to examine only top-scoring genomic neighborhoods based on sequence similarity to a query protein.

The interactive browser-component of AnnoView (Figure 1) allows users to explore multiple gene neighborhoods (up to 500) in a single window. A toolbar panel at the top of the browser allows users to perform different functions. These include a zoom-in and zoom-out function, and a download button for retrieving gene neighborhood information in CSV format. Downloaded CSV data can be customized by the user to edit existing gene neighborhood information or add taxonomic and functional annotations. The gene neighborhood plot can also be exported in vector-quality SVG format suitable for publication or offline editing by image-editing software (e.g., Adobe Illustrator, Inkscape, Affinity Designer).

**Figure 1.**
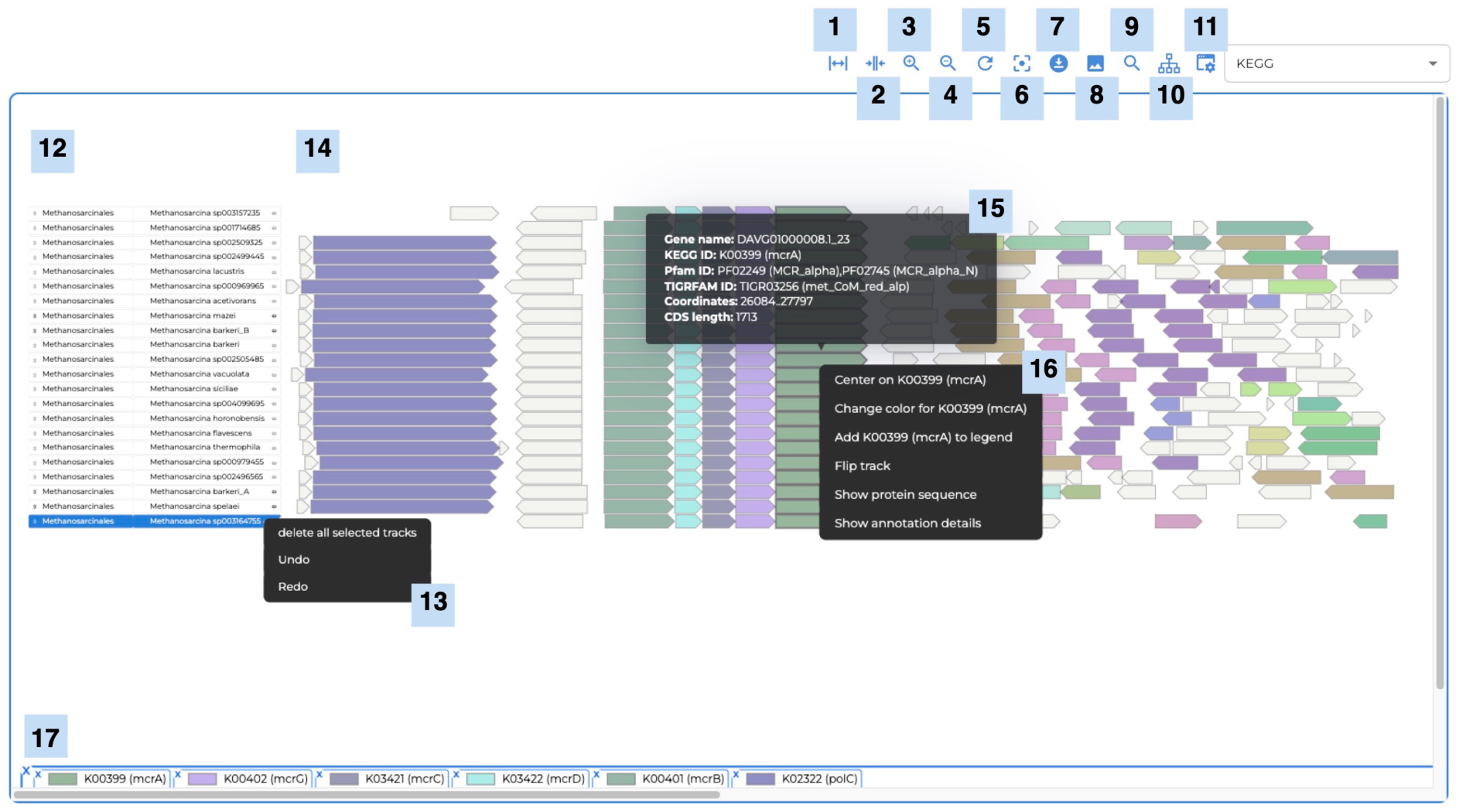
The AnnoView gene neighborhood browser and its key features. (1) Stretch; (2) Compress; (3) Zoom in; (4) Zoom out; (5) Reset the page; (6) Recenter image; (7) Download the gene neighborhood in CSV format; (8) Export image in SVG format; (9) Search box; (10) Display taxonomy labels; (11) Color genes by metadata; the default is colored by KEGG annotations; (12) Taxonomy labels; (13) Delete selected tracks; (14) Gene neighborhood; (15) Gene information when hovering on a gene; (16) More options when right-clicking on a gene; (17) Legend.

To navigate to a specific gene, users can search for gene names in the search box. The search box is also useful for navigating genes that share a general term such as “flagellar” to highlight multiple matches (e.g., “*flagellar* M-ring protein FliF” and “*flagellar* motor switch protein FliG”). Taxonomy labels allow users to choose the taxonomy level to display in the visualization panel. Users can change the displayed metadata to color gene neighborhoods based on different functional annotations.

The visualization panel is an interactive interface that allows exploration of basic gene information, clustering of gene neighborhoods, editing of the plot, and data retrievals, such as protein sequences and gene metadata (**Figure 1**). For each gene neighborhood displayed in the visualization panel, its taxonomy information is displayed on the left side of the panel. The taxonomic level can be changed under the taxonomy labels in the toolbar panel. To view gene information, the user can either hover over a gene or right-click on a gene for more information (e.g., protein sequence and annotation details). More options are available when right-clicking on a gene for the user to edit the gene neighborhood plot in AnnoView. After choosing a center gene, gene neighborhoods can be clustered using the clustering algorithm implemented in AnnoView, which is based on the similarity of neighboring gene content (see Methods, **Supplementary Figure 2**). Furthermore, users can flip gene neighborhood orientations, change gene colours, and add genes to the legend. The vertical order of gene neighborhoods displayed can also be changed based on a user’s preference. For example, the user may wish to manually re-order neighborhoods based on phylogenetic relationships. A unique token is generated for each online session and can be used for access for up to a month. Because of the unique web token, the URL for the user’s current session can also be shared with other users over the web to enable data sharing.

## Example Workflows

### Example 1: Using AnnoTree-DB search to explore the McrA operon

We envision that a common use case for AnnoView will be for users to search our pre-computed genome database for matches to a gene/protein of interest. For instance, using AnnoView, a user may explore gene neighborhoods surrounding methyl-coenzyme M reductase subunit alpha (McrA) in archaea (**Figure 2**). Methyl-coenzyme M reductase (MCR) is an enzyme that catalyzes the final step of methanogenesis in archaea [54]. Searching a representative McrA protein sequence from *Methanosarcina barkeri* (https://uniprot.org/uniprotkb/P07962/) in the AnnoView database returns 379 detected homologs. As mentioned in the previous section, there are two ways of filtering the database matches: either select hits by taxonomy or by the top N (where N is defined by the user) number of hits ranked by *E*-value. Here, we select McrA homologous sequences in *Methanosarcina*, a genus of archaea known for methanogenesis. The gene neighborhood visualization shows that *mcrA* is part of a highly conserved *mcrBDCGA* gene cluster, which forms the known operon structure of methanogenic MCRs [55]. The MCR operon consists of the essential genes *mcrA*, *mcrB*, and *mcrG*, as well as two accessory genes *mcrC* and *mcrD*.

**Figure 2.**
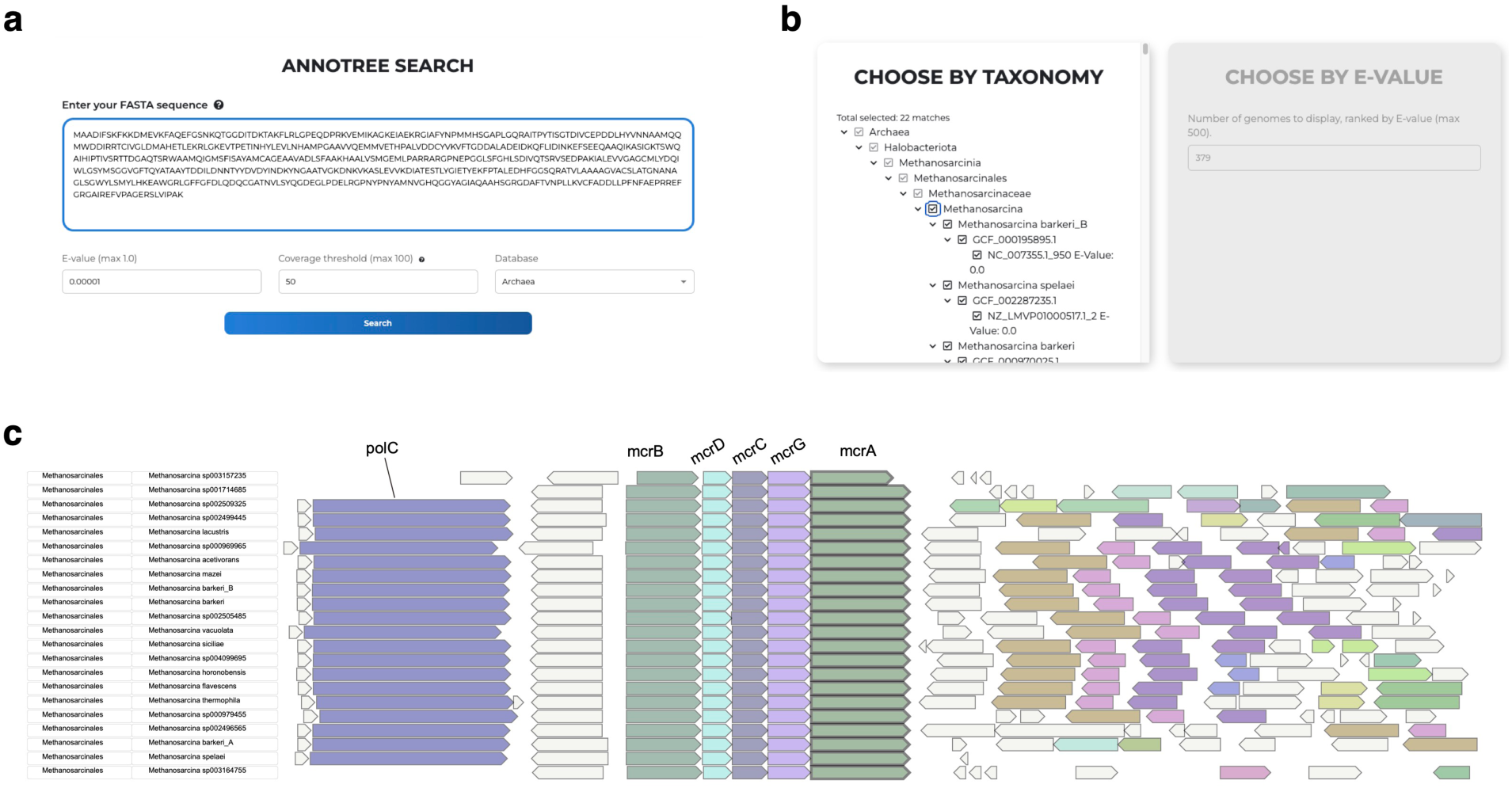
An example illustrating an AnnoTree database search using AnnoView. (**a**) AnnoTree search interface. (**b**) The intermediate page for user to select gene neighborhood by taxonomy or by *E*-value. (**c**) The genomic context of selected McrA homologs in the *Methanosarcina* genus.

### Example 2: Visualizing a custom gene neighborhood dataset (the SLR4 gene locus)

Another envisioned use case for AnnoView is to upload a custom gene neighborhood dataset and explore it further using AnnoView’s gene neighborhood browser (**Figure 3a,b**). The gene neighborhood dataset can be generated from public genomic databases such as NCBI or user-generated genomic datasets in GBK or GFF format. To visualize the gene neighborhoods labeled by different functional annotations than those provided by GenBank (e.g., protein product name), such as Pfam or KEGG annotation, a bioinformatic workflow is provided at https://github.com/satellite830/AnnoView. This workflow outlines how users can download a CSV table from AnnoView, add customized annotations and taxonomy information to the table, and upload the data back to AnnoView for further exploration (See Methods for a detailed workflow).

**Figure 3.**
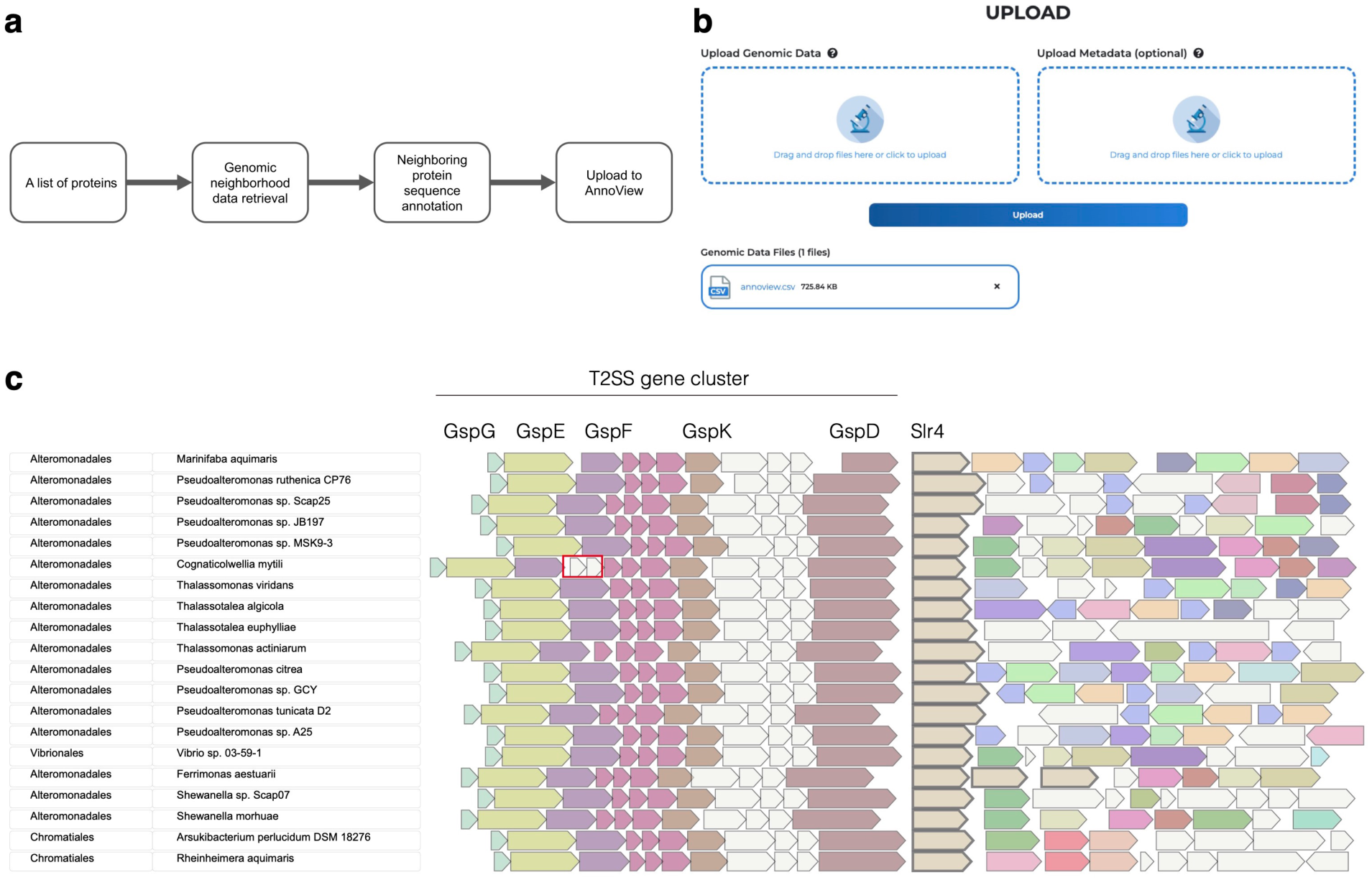
AnnoView exploration of a custom uploaded gene neighborhood dataset. (**a**) Workflow of generating the gene neighborhood dataset. The gene neighborhood dataset in GBK format was downloaded from NCBI, then the neighboring genes were functionally annotated by Pfam. The gene neighborhood dataset along with its protein functional annotations was uploaded to AnnoView for visualization. (**b**) AnnoView upload interface. (**c**) The gene neighborhoods of Slr4 and its homologs. The species-specific insertion in *C. mytili* is highlighted as a red box.

To demonstrate this workflow, we focus on an example from a previous publication [56], where AnnoView was used to visualize gene neighborhoods surrounding a newly identified protein called Slr4 from *Pseudoalteromonas tunicata* and its homologs in related species (**Figure 3c**). Gene neighborhoods were annotated and visualized based on Pfam annotations and all annotations occurring more than twice have been highlighted (**Figure 3c**). In this study, Slr4 was functionally characterized as a novel surface layer (S-layer) protein that forms a proteinaceous protective layer surrounding cells [56]. The genomic context provides additional supportive evidence of this function, as it reveals a conserved pattern of Type II secretion (T2SS) pathway genes immediately upstream of Slr4, which have been previously implicated in S-layer protein secretion (**Figure 3c**). Because dedicated T2SS systems are a common feature of S-layer genes [57, 58], it is likely that this conserved operon upstream of *Slr4* plays a similar function and serves as a dedicated T2SS pathway for secreting this novel S-layer protein family.

In addition to visualizing conserved neighborhood features, another important function of AnnoView visualization is to reveal differences in genomic context. An example of this is a species-specific genomic insertion in the Slr4 neighborhood of *Cognaticolwellia mytili* (**Figure 3c**). The insertion includes two additional ORFs annotated as PF18765 (polymerase beta, nucleotidyltransferase) and PF01934 (ribonuclease HepT-like), which are two genes in the recently discovered type II toxin/antitoxin (HEPN/MNT) system [59]. The insertion of this toxin/antitoxin system into the T2SS gene cluster is consistent with the highly mobile nature of type II toxin/antitoxin systems [60].

### Example 3: Visual comparison of Betacoronavirus genomes

While primarily intended as a tool to explore microbial gene neighborhoods around genes of interest, AnnoView is also capable of whole genome visualization and comparative analysis for relatively smaller sequences, such as viral genomes, plasmids, and in some cases, smaller bacterial genomes. An AnnoView visualization of several *Betacoronavirus* genomes reveals several interesting and biologically relevant differences in genome structure (**Figure 4**). The overall genome structure (gene order and composition) of different *Betacoronavirus* species is highly conserved. Important genes such as ORF1ab, N (nucleocapsid protein), S (spike protein), M (membrane protein), and E (envelope protein) are conserved in all *Betacoronavirus* genomes. However, HE (hemagglutinin esterase) is only present in the *Embecovirus* subgroup. The hemagglutinin esterase is an enzyme that facilitates attachment and destruction of sialic acid receptors present on the host cell surface, which aids in molecular recognition of specific target cells [61, 62]. The unique insertion of the HE gene into genomes within the *Embecovirus* lineage likely resulted from a horizontal transfer event originating from influenza C/D [63]. This proposed insertion event likely occurred in ancestral members of the *Embecovirus* subgenus before the divergence of other *Betacoronavirus* subgroups, resulting in the unique presence of the HE protein in the *Embecovirus* subgroup.

**Figure 4.**
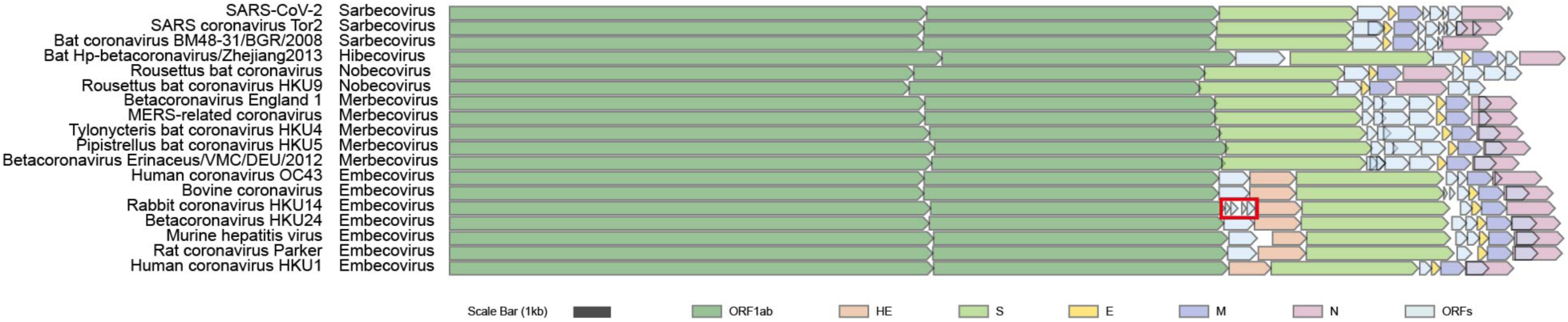
Visualization of *Betacoronavirus* genomes using AnnoView. Genome tracks are sorted by the genome phylogeny from NCBI Virus. The red box highlights a unique split gene in the rabbit coronavirus HKU14 genome.

A second genomic difference revealed by AnnoView visualization is the presence of four ORFs between the ORF1ab and downstream HE gene within the rabbit coronavirus HKU14, in comparison to other genomes from the *Embecovirus* subgroup, which have zero or one ORF (NS2a) between ORF1ab and HE (**Figure 4**). Previous work has shown that this unique pattern in the rabbit coronavirus genome have resulted from the splitting of an ancestral NS2a gene into four smaller ORFs [64].

## Conclusion

AnnoView is a versatile gene neighborhood visualization tool that enables rapid and user-friendly exploration of microbial genomes. The AnnoView database is integrated with AnnoTree, which provides comprehensive and consistent taxonomic classification for all genomes within the Genome Taxonomy Database. In addition, users can explore their own user-uploaded genomic datasets, including orthologous gene neighborhoods across sets of species or small size genomes. AnnoView has already enabled genomic analysis and visualization for several studies [7, 56, 65–67]. In the future, we hope to add more features to AnnoView, such as expanding the set of functional annotation methods available to users (e.g., including the InterPro suite of annotations [68]), updating AnnoView on a regular basis with the most up-to-date GTDB data, allowing users to customize their genome neighborhood criteria (e.g., window size), and displaying other genomic features, such as tRNAs, rRNAs, CRISPR loci, and pseudogenes. We envision AnnoView being a useful tool for gene neighborhood exploration, visualization, and comparison of microbial genomes with applications for genetics and genomics research more broadly.

---

**Supplementary Figure 1.**
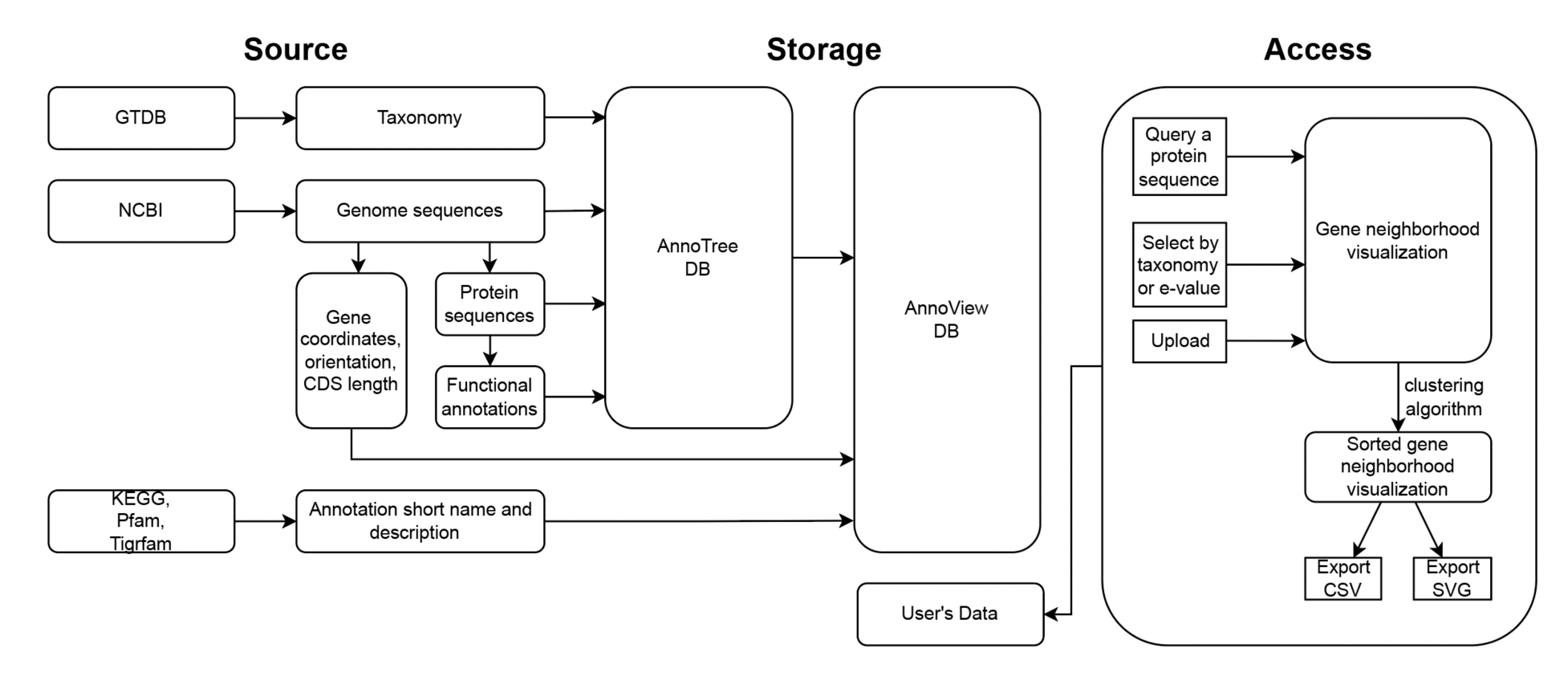
Software architecture diagram showing the components of AnnoView.

**Supplementary Figure 2.**
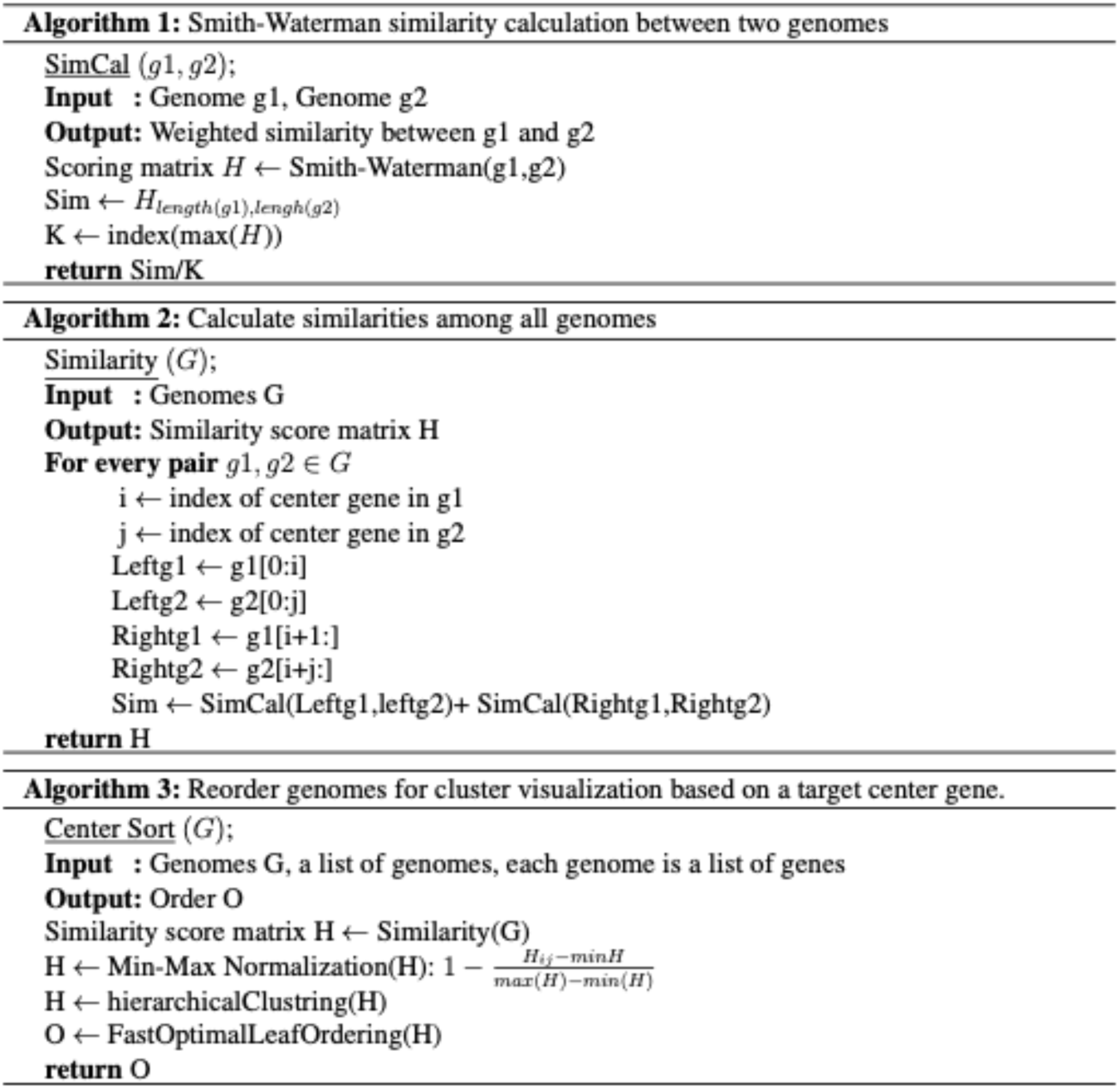
Pseudocode for the clustering algorithm that sorts gene neighborhoods based on a target center gene.

**Supplementary Figure 3.**
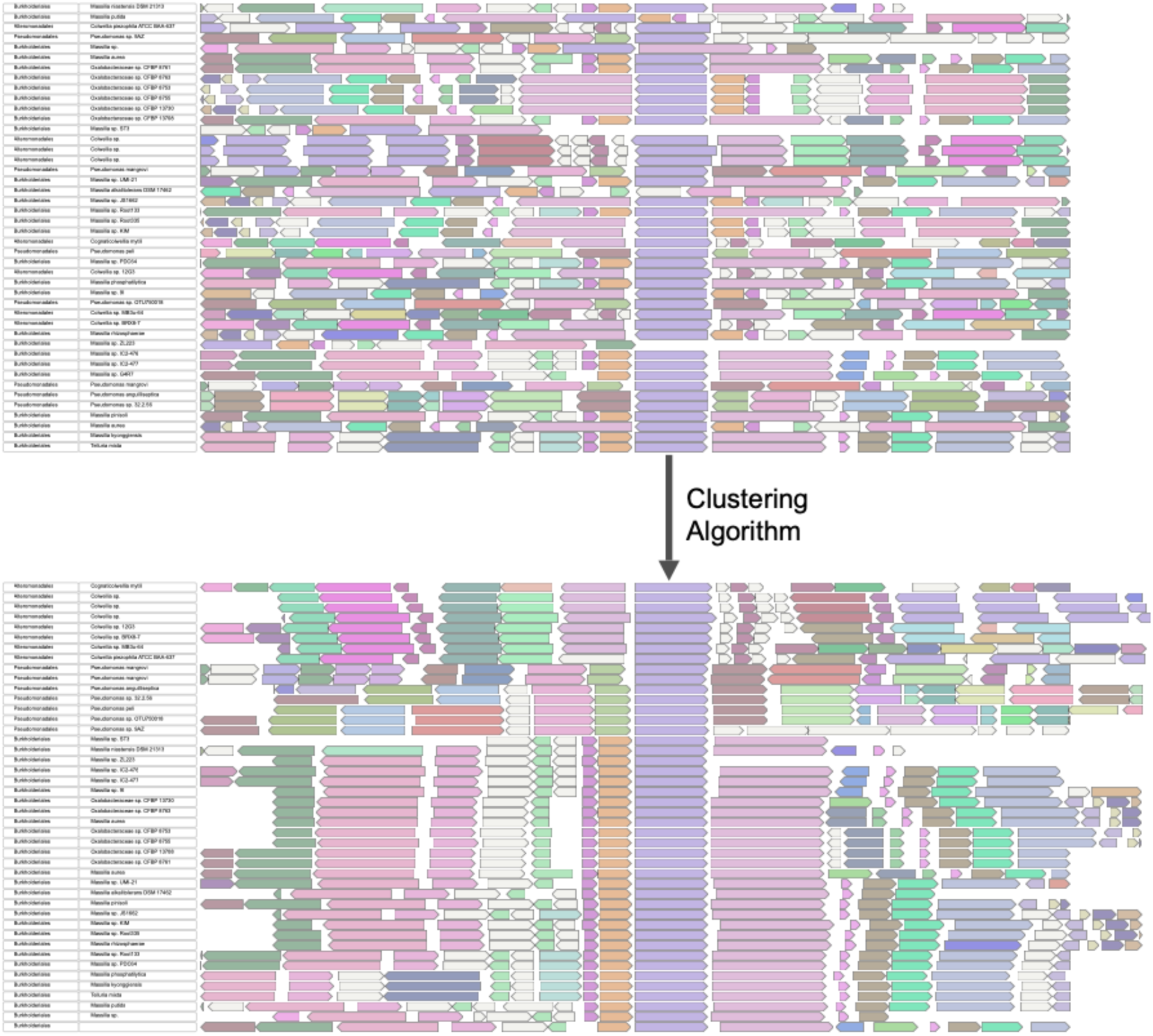
An example demonstrating the gene neighborhood clustering algorithm. A pre-clustered set of gene neighborhoods is shown above. The neighborhoods are then clustered and sorted (below) based on flagellinolysin [69] selected as the center gene (purple). Gene neighborhoods are clustered based on gene composition, which is defined based on their lists of functional annotations. After sorting, patterns emerge which illustrate evolutionary groupings of genomic regions based on their common ancestry.

